# FloraTraiter: Automated parsing of traits from descriptive biodiversity literature

**DOI:** 10.1101/2023.06.06.543883

**Authors:** R.A. Folk, R.P. Guralnick, R.T. LaFrance

## Abstract

*Premise—*Plant trait data are essential for quantifying biodiversity and function across Earth, but these data are challenging to acquire for large studies. Diverse strategies are needed, including the liberation of heritage data locked within specialist literature such as floras and taxonomic monographs. Here we report FloraTraiter, a novel approach using rule-based natural language processing (NLP) to parse computable trait data from biodiversity literature.

*Methods and Results—*FloraTraiter was implemented through collaborative work between programmers and botanical experts, and customized for both online floras and scanned literature. We report a strategy spanning OCR, recognition of taxa, iterative building of traits, and establishing linkages among all of these, as well as curational tools and code for turning these results into standard morphological matrices. Over 95% of treatment content was successfully parsed for traits with < 1% error. Data for more than 700 taxa are reported including a demonstration of common downstream uses.

*Conclusions—*We identify strategies, applications, tips, and challenges that we hope will facilitate future similar efforts to produce large open-source trait datasets for broad community reuse. Largely automated tools like FloraTraiter will be an important addition to the toolkit for assembling trait data at scale.

---

Botanists have been gathering information on plant traits, which comprise the entirety of measurable aspects of plant phenotypes, since the dawn of scientific pursuit (Fig. 1a). With traits scored for many species, biologists can ask questions about phenotypic and functional differences spanning the plant tree of life, and within and across ecological communities. These questions are fundamental because traits underlie how species interact with and adapt to their surroundings and how we as humans interact with them (Freudenstein et al., 2016). Yet despite the long history of studying plant traits, this information remains mostly unavailable in a form that is usable for quantitative analysis (Hortal et al., 2015). A shortage of “computable” trait data has a very real effect on our view of global plant diversity (Pakeman and Quested, 2007; Pakeman, 2014; Sandel et al., 2015; Cornwell et al., 2019), and tends to be much higher in quantitative terms than other plant data and information domains such as DNA and geographic data (Sandel et al., 2015; Folk et al., 2018). This lack is distributed along biogeographic, socioeconomic, political, and other axes that impact the science performed in regions of Earth where this same information is urgently needed (Meyer et al., 2015; Daru et al., 2018; Cornwell et al., 2019).

**Fig. 1.**
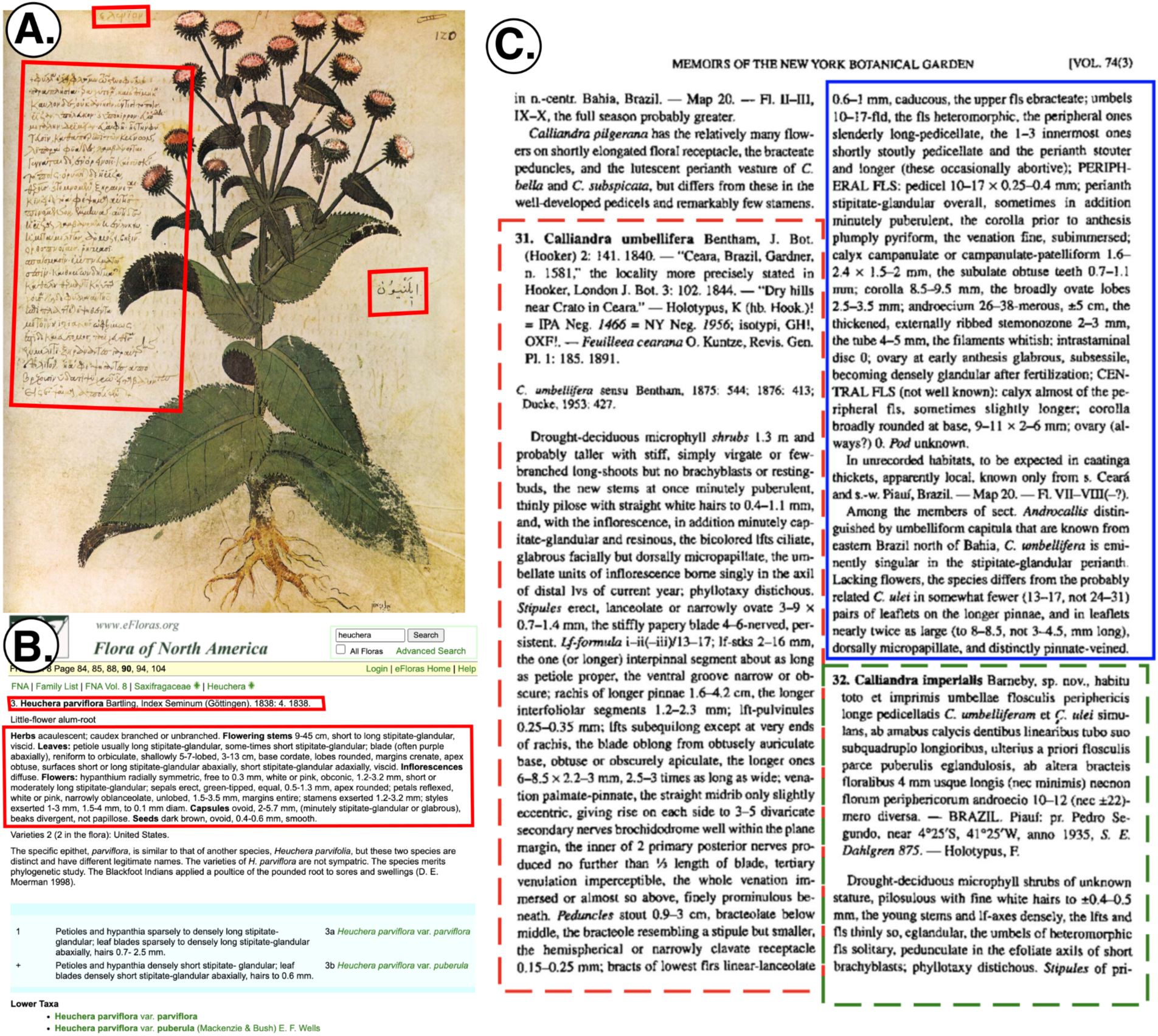
Anatomy of a piece of biodiversity literature. (A) The collection of plant trait information, while a modern challenge, has a long history. The *Vienna Dioscorides*, a manuscript from the early 6th century, represents the earliest direct and relatively complete survival of scientific botanical descriptions in the Western tradition and illustrates the long history of study that underlies our current understanding of plant traits. The structure of the entry for *Inula helenium* L. follows many aspects of a modern description and therefore illustrates concepts, with its taxon names in Greek (έλενίου, marked in red, upper left) and Arabic (ِﺳﻦ َرا, marked in red, right) marked, as well as the original description (marked in red, lower left) including distribution, morphology, habitat, and medical uses in that order. Our purpose here is to break down the latter (the *description*) to basic *traits* and link these to the former (the *taxon*). (B) Entry for *Heuchera parviflora* from *Flora of North America*, a more modern e-flora example, where the text is already machine-readable and relevant portions of the description structure can be fetched by following links and cleaning up HTML tags. (C) A more difficult use case from a monographic work treating *Calliandra*: while also a recent treatment, this is a scanned text that had to be subjected to OCR (optical character recognition) according to the segmentation strategy reported in Methods to generate machine-readable text. Structural features like page breaks and page headers are not description or taxon text and therefore should be removed from OCR; this was done *a priori* here with the boxes marking the start of the first full treatment on the page; the blue box continues this treatment and the green box indicates a second treatment (this is a screenshot of one of the processing tools presented here; see Methods). All of these text boxes had to be annealed correctly before further processing. Also to be seen are various formatting peculiarities (capitalization, italicization, font size, indentation, unusual abbreviations, multilingual material, some of which parallel [A]) that vary by work and can be variously complex or inconsistent; including or discarding these is the subject of optimizing particular sources.

If plant traits are needed and missing, which traits should we measure and how do we best measure them? As argued by Violle et al. (2014) and Hortal et al. (2015), “the traits that are generally measured are often the most simple, rather than the most functional” (Hortal et al., 2015 p. 529) whereas many ecologists might prefer the traits most strongly linked to ecosystem function be measured first. But rather than ask which traits are best measured anew, it is equally important to assure that we best leverage the vast information we already have. Information about many traits measured across the entire plant body for many species has long been disseminated by specialists, but these data are locked up in the inaccessible form of biodiversity literature (Rinaldo, 2009; Thessen et al., 2012; Folk et al., 2018; Penev et al., 2019; Folk and Siniscalchi, 2021; Shirey et al., 2022). For our purposes, “biodiversity literature” refers to all forms of scientific and paraprofessional output that contains summary statements regarding *taxa* and their *traits*. This includes species descriptions, floras, field guides, monographic revisions, and similar works (examples in Fig. 1). Such biodiversity literature is unique in that, rather than representing a directly verifiable measurement on a single organism, these resources provide measurements that represent an expert’s judgment regarding a set of observations contingent on a taxon hypothesis. A literature-derived trait therefore has the downside that it is only as good as the taxonomic delimitation that underlies it. This same property is a key strength: unlike other sources of data, biodiversity literature represents the assessment of a domain expert, such as a botanist specializing in a family of plants, and therefore likely to represent the state of the art regarding currently understood taxon boundaries and attributes. An expert is able to identify and exclude diseased and underdeveloped plant organs, can interpret the sometimes complex structural homology between plant species, and characteristically will focus on the structures most variable among closely related organisms. Biodiversity literature extraction therefore holds promise as a major source of plant trait information and a complement to other sources, but despite several previous proof-of-concept reports (Cui et al., 2016; Endara et al., 2018) the botanical community still lacks a general-purpose approach that can generate high-quality data for many species.

In this paper, we report on work that far extends pipelines originally designed for vertebrate measurements (dubbed Traiter in Guralnick et al., 2016) to extract morphological data from floristic publications. Traiter originally used regular expressions to process text data about body length and mass from vertebrate specimen records and produce harmonized, standardized quantitative measurements.

However, a regex approach has some important downsides when applied to unstructured publication text. First, it relies on text data sources that are already highly structured, such as specimen labels with free text entry for body size, embryo counts, and similar content. While there are structured reports of traits in flora or other resources, much of the data is in prose not presorted into trait categories, which often leads to extensive human effort curating the resulting parses. Second, regexes rely on patterns of characters, making it difficult to use other information not contained in the characters themselves, such as that implied by parts of speech or sentence structure. For instance, consider the description of *Comptonia peregrina* from *Flora of North America*, according to which the leaves are “3-15.5 × 0.3-2.9 cm, lobes alternate to nearly opposite, base truncate, cuneate to attenuate, or oblique, apex acute; surfaces abaxially pale gray-green, densely pilose to puberulent, adaxially dark green, densely pilose to glabrate, gland-dotted, especially adaxially.” To correctly understand the information in this description requires recognizing several things; 1) that every structure mentioned is a subpart of a leaf (our dataset must therefore represent or at least be cognizant of hierarchical structure); and 2) That the given measurements go with the entire leaf but all other descriptors belong to subparts (we cannot rely on word order even though the adjectives follow the nouns here, according to a typical regex strategy; this varies between publications or even between sentences). In both cases, we must use context to understand that the first reported numbers represent length and the second are width, and we must navigate qualifiers (“especially”, “nearly”), positional information (“adaxially”), and multiple measurements with complex relational prepositions and conjunctions (“cuneate to attenuate, or oblique”). In short, a flexible method to extract data from text like this must be able to understand the structure of a typical sentence from a taxon description.

Here we report a new approach called FloraTraiter that shifts to using natural language processing (NLP) approach to more flexibly extract traits. The specific form of NLP used here, rule-based parsing, leverages preexisting language models to break down biodiversity descriptions into parts of speech, with an extended vocabulary to handle technical botanical descriptions. Then a series of newly implemented steps further process structural elements that pertain specifically to a biodiversity description, beginning with recognizing a taxon and then using an interactive process to identify partial traits and map them to taxa. We test the new approach on descriptions of the plant order Fagales as reported in online *eFloras* treatments and implement code to translate parsed descriptions into a recognizable morphological matrix format. We then score trait extractions against human observers examining the same source references to count false negatives and false positives, giving us a quantification of traits that are missed or incorrectly scored respectively. Finally, we demonstrate some standard downstream uses for the trait extractions.

## METHODS

*Raw data—*Parsing begins with literature that has either already been entered digitally or is an image file that must be subjected to OCR (optical character recognition). OCR is the most complex task and so this will be described first and at length. Commercial solutions for PDF OCR are available, such as Abbyy FineReader, but a custom approach was used because standard OCR packages were found unsuitable after extensive testing. Problems with standard OCR include (1) OCR text, while appropriately placed on the page, is commonly not in reading order when extracted; (2) sometimes text on the page is missing; (3) frequently characters are substituted for other similar-looking characters (such as exchanges involving u, n, m); and most importantly (4) many aspects of page structure are a nuisance irrelevant to the data contained in a page; these aspects increase the challenge of parsing traits, and especially linking them back to taxa when descriptions span pages. Points 3 and 4 will be discussed further below.

The first step is to convert the PDF into images, one image per page. This function is captured in “pdf_to_images.py” (hereafter all quoted scripts are found at: https://github.com/rafelafrance/FloraTraiter), which is a wrapper around the pdftpcairo program, a module in poppler-utils (pre-compiled command-line programs from the Poppler library; https://poppler.freedesktop.org/) to be installed separately. Second, the PDF images are manually segmented by drawing bounding boxes around text in reading order and marking which ones denote the start of a treatment; these bounding boxes indicate where each treatment starts and ends while marking the order of the treatment text. This is done in a Python script called “slice.py”. The colors help indicate the order in which a page is read, with red always denoting the first box, blue the second, etc.; the dashed box outlines indicate a treatment start (Fig. 1c shows an actual session using this program). Only treatment text is outlined with figures, captions, headers, and similar materials left out. This is essential because large amounts of structural information in text is a nuisance and must be discarded, and the flow of text on the page is not always easily determined programmatically. Nuisance text includes page breaks and number breaks, as well as material that does not contain formal descriptions, such as literature surveys and indices. Handling this cleanly is important because successfully identifying breaks in descriptions is essential for discovering links between traits and the taxa they belong to.

Once the PDF images are segmented, OCR is conducted on the text in each bounding box; boxes of text are then stitched together with markers in the text to indicate the start of treatments. This script is called “stitch.py”. Finally, common OCR errors are corrected and the text is normalized with “clean_text.py”. As a corollary, limited typo correction can be handled but in general high-quality and well-aligned scans are needed to produce good OCR content.

The raw data for HTML sources is much simpler, and involves “spidering” (iteratively traversing the structure of) a base webpage for a taxon and pulling taxonomic treatments guided by HTML markup. Code for this purpose is also available at https://github.com/rafelafrance/FloraTraiter.

*Controlled vocabulary—*The approach described in the following section relies on pre-built language models, but these are generally trained on text intended for a wide audience and lack vocabulary on specialist topics. Conversations between botanists and programmers on this project led to us identifying a basic botanical trait vocabulary to add to the model; early drafts were distributed to organismal specialists for comment, which led to identification of missing technical words (the final vocabulary developed is integrated throughout the GitHub repository in relevant processing steps).

Fortunately, differing sources on similar plants tend to differ little in vocabulary, with the main need being to identify several commonly used synonyms and closely related terms that reflect differing authorial habits and editorial policies (pod = legume, androecium ∼ stamens). The greater effort in adapting code to new projects has been in shifting taxa, as highly specialized plant families will have numerous trait terms that are not of broad application or may have restricted meanings in context.

We also built a custom vocabulary for identifying scientific names within text. This comprises a combination of four sources of known binomials and monomials assembled from these sources: ITIS (SQLite version at https://www.itis.gov/downloads/index.html), The WFO Plant List (https://wfoplantlist.org/plant-list/classifications), Plant of the World Online (http://sftp.kew.org/pub/data-<u>repositories/WCVP/</u>), and further miscellaneous taxa not found in the other sources (https://github.com/rafelafrance/FloraTraiter/blob/main/flora/pylib/traits/terms/other_taxa.csv)

*Parsing strategy—*The next major step is to parse each treatment in the resulting text. This uses a wrapper script called “extract_traits.py” (hereafter, all quoted scripts are from the base repository at https://github.com/rafelafrance/FloraTraiter unless otherwise specified) to call code from the general Traiter repository (https://github.com/rafelafrance/traiter). These two repositories use a combination of rules and statistical methods to parse traits in the treatments. The basic parsing approach is a hybrid statistical and rule-based approach relying heavily on the spaCy Natural Language Processing (NLP) Python library. spaCy is used for NLP because of its prebuilt statistical models and its flexible framework for building parsing rules custom to this project. First, spaCy’s statistical models are used for determining the parts of speech (POS) to which each word belongs. POS is used when building rules for parsing. For instance, parsing a full taxon can be approached by looking for a binomial (species) notation followed by a proper noun or a set of proper nouns (like: John Smythe and Jane Jones) separated by conjunctions to find candidate taxon authorities. We also take advantage of customizability in spaCy: for instance, its default tokenizer (breaking texting into meaningful words or word parts) is intended for general-use text such as Wikipedia but tends to fail in tokenizing the formalized and idiosyncratic text of taxonomic treatments. Addition of custom vocabularies greatly improves performance in this respect.

A full outline of the parsing pipeline (summarized and simplified in Fig. 2) proceeds as follows: (1) Use a customized tokenizer to break treatment text into tokens. (2) Allow a spaCy model to identify what POS each token belongs to as well as other things like its lemma (the “normalized” version of a token that reduces it to its root meaning; e.g., “good” would be the lemma of “better”). (3) Parse traits using rules and phrases. Most traits have a predefined vocabulary of words and phrases (noted above) that anchor rules for further parsing. These terms are obtained from organismal experts and other authoritative sources; as described above, the approach used for this study consisted of iterative improvements of drafts that were submitted to expert botanists for comment. The initial product of parsing is termed a “partial trait.” (4) Use the anchor phrases in rules that build up full traits from partial traits. This involves finding patterns of words and symbols around anchors identified already to build up the traits themselves.

**Fig. 2.**
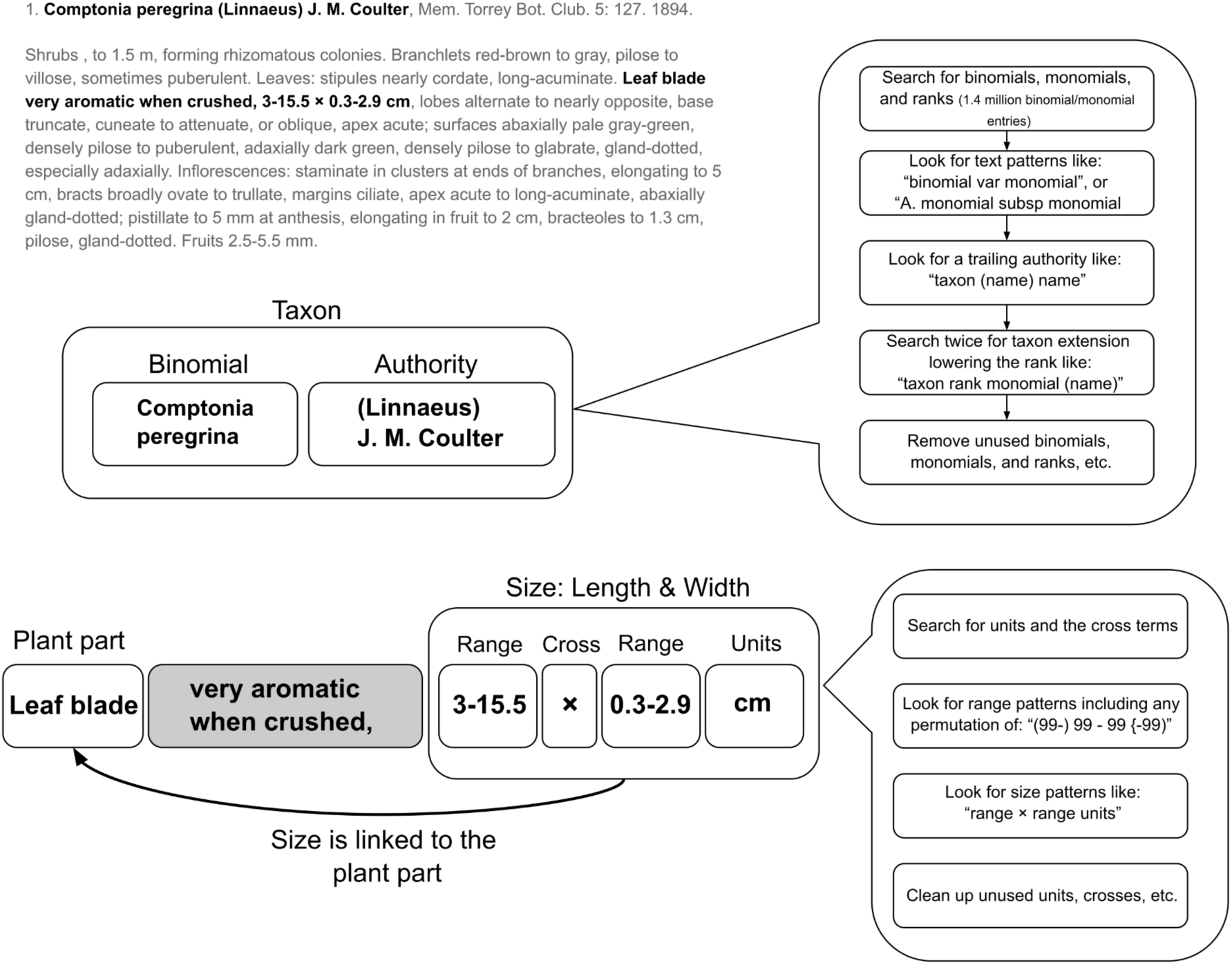
Workflow summary of FloraTraiter. Taking the example from the introduction (top left), in bold is the taxon and the first part of the leaf description of *Comptonia peregrina* in *Flora of North America.* The first step is to identify a “taxon anchor” (center) which involves finding binomials or monomials in the text, and searching the vicinity for additional material representing authors and subspecific names and ranks. Once this taxon anchor has been identified, plant parts are identified and linked to the taxon (bottom; shown is a measurement example, where the process begins with a similar anchoring process around units and similar symbology) and finally attributes such as measurements are linked to them to yield “partial traits.” The given example demonstrates the frequent use of context clues such as implied length and width, as well as the common need to link text material that is non-adjacent.

Sometimes these phrases are the final trait, but more often this step is repeated to build up larger and larger traits, where order of steps matters and is controlled in repeated iterations to build traits into their final form. (5) Clean up any leftover phrases and partially applied rules so the words/tokens can be used in other traits. For instance, in Fig. 2, information about leaf scent (grayed out) splices a statement about leaf length and width, but this discarded information can be used to populate a trait about general leaf properties. (6) Link traits after they are parsed. All built-up traits are linked to the taxon in the treatment title, and other traits like plant or flower parts get linked based on proximity and other other trait-specific considerations like rules about whether links across sentences are allowed. Rules are also applied based on the legal structure of linkages considering the nature of the trait: a plant part may have several colors but typically only has one size, barring sexual dimorphism; multiple entries are forbidden in certain contexts.

In general the overall process follows this outline, but special strategies were also used. Some traits can be ambiguous because their meaning depends on context: for instance, Green can be a last name, a color, or an administrative unit, so context is used extensively in parsing to distinguish among related meanings.

*Curation—*An important component of achieving high-quality extractions was a series of structured conversations between botanists and programmers as the project developed. This was facilitated by the preparation of interactive visuals that enable non-specialists to understand how the NLP method is reading meaning into and pulling information out of text. An example of marked up HTML output may be seen here: https://htmlpreview.github.io/?https://github.com/ryanafolk/fagales_traits/blob/main/trait_processing/Fagales_2023-01-23/Fagales_2023-01-23.html (Fagales content for *Flora of North America*). Each color corresponds to a trait, with trait labels corresponding to CSV outputs (below). Common colors between highlighted sections of the treatment and extracted trait data speed up the process of comparing the two, as do different options for color-coding traits. Special data models for reported traits apply to complex numerical data as reported in standard botanical descriptions. For instance, “(1-)2-4(–5) cm” is separately parsed into measurements labeled “min,” “low,” “high,” and “max,” as well as a field representing measurement units. HTML outputs were distributed and marked up by participating botanists for correction in the form of prose commentary or marked-up CSV outputs in several iterations.

*Post-processing—*The output of FloraTraiter, even after extensive curation, looks very different from a standard morphological matrix (Fig. 3a). Because the NLP process aims to capture as much text as possible, (1) numerous fields are returned because species descriptions discuss numerous structures. Parts of the internal data structure also lead to this, as “flattened” CSV outputs use separate fields for measurement ranges and multiple values. Additionally, (2) while basic botanical vocabularies are integrated, the philosophy employed here was conservative in the matter of trait synonymy and generally measurements and descriptions were kept separate that could conceivably be different (seed width and height are separate even if only one is mentioned, fruiting and flowering pedicel measurements are not combined, stamen and androecium length were not considered identical, etc.). Finally, (3) integrating a large taxonomic and literature scope leads to large, sparse matrices because different groups have some commonalities but numerous differences that reflect specialized morphology. Because a typical downstream user will likely have a biologist’s training, we felt it was important to simulate a data structure closer to a morphological matrix.

**Fig. 3.**
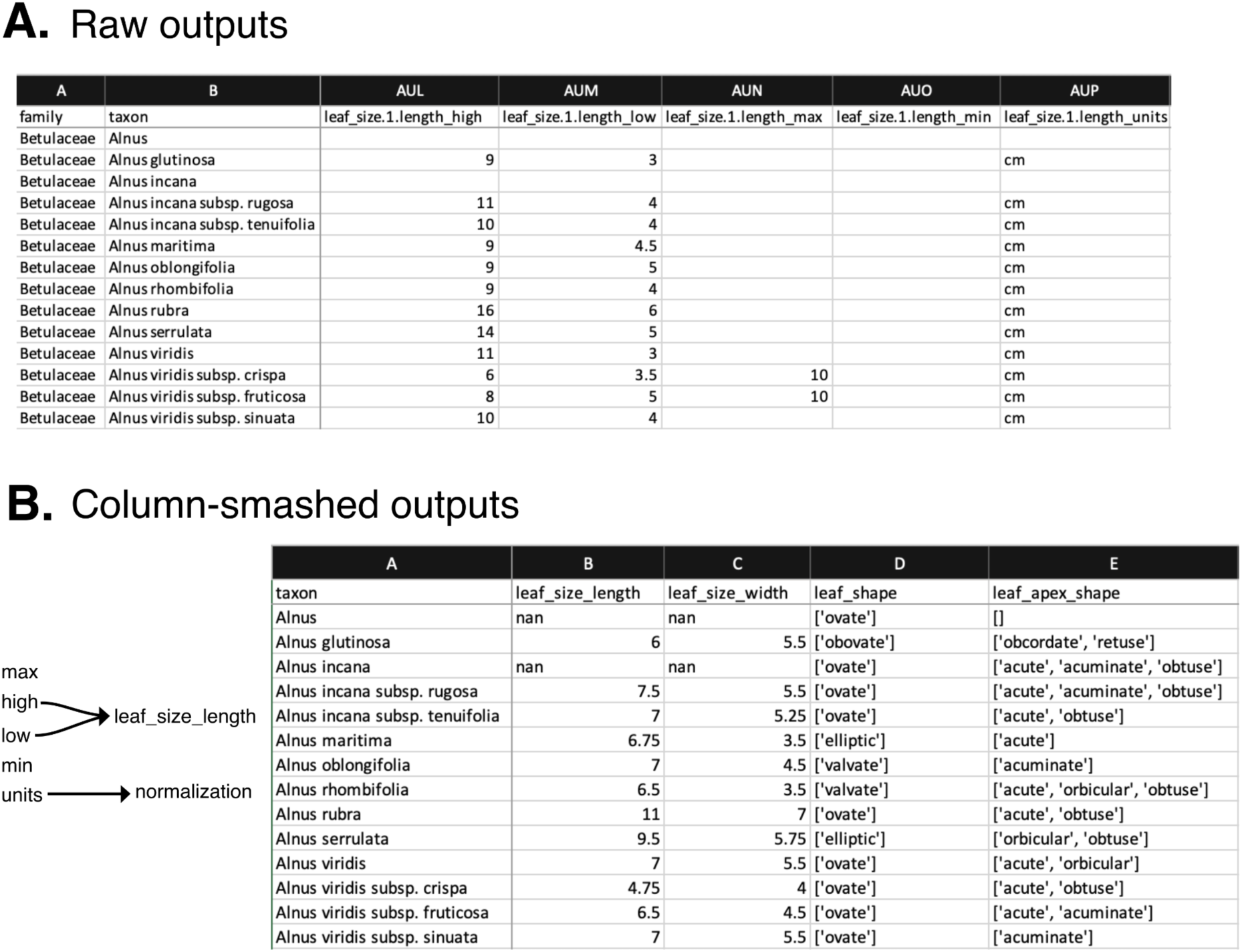
(A) Raw FloraTraiter output, demonstrating the very long field format due to the detailed method of parsing traits. (B) A result of “column-smashing,” where quantitative data were reported as the midpoint excluding extreme measurements, and qualitative data were reported as a list.

**Fig. 4.**
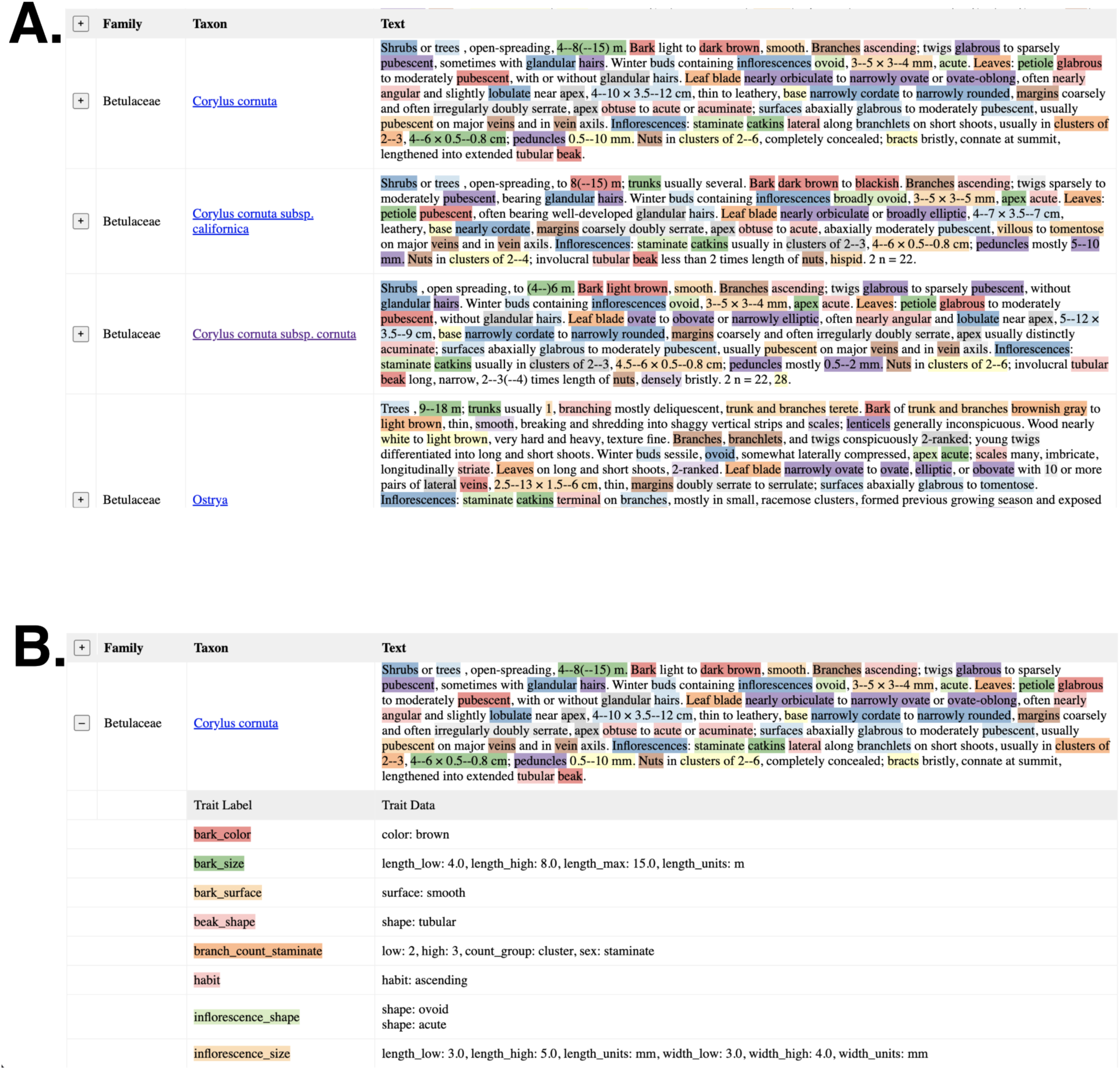
Demonstration of the HTML outputs. Two screenshots are shown for *Corylus* from *Flora of North America*; (A) shows the unexpanded view (the “+” symbol on the HTML will expand the view and show the atomized traits as shown in [B]), with every highlighted portion corresponding to trait content that was extracted. (B) shows an expanded view of one species with the actual scores for each trait. This view allows organismal experts to evaluate the operation of FloraTraiter’s NLP approach.

The approach implemented is a generalization of the morphological matrix harmonization reported in Folk et al. (2019), deposited on GitHub at https://github.com/ryanafolk/fagales_traits/. First, a controlled vocabulary for fields is specified by a simple spreadsheet format. This is easily edited by a non-specialist to map field definitions that can be combined. Fields are specified by three separate CSVs to represent (1) categorical data that should be represented as a list, termed “concatenate_terms.csv”; (2) count data that should be summarized in ways appropriate for cardinal numbers such as the mode, termed “range_terms_count.csv”; and (3) quantitative measurements that can be summarized by the mean or other methods appropriate for continuous data, termed “range_terms_quantitative.csv”. Additional controlled vocabularies specify (4) equivalency among sex terms (e.g., male and staminate flowers are the same) in “sexes.csv”, (5) terms to be excluded in “discard_terms.csv”, and (6) measurement unit data in “unit_columns.csv”. Second, a synonymy among data in the fields is established in “synonyms.csv”. This data file represents some of the most “opinionated” decisions as it includes judgements about term usage: things that certainly mean the same thing such as “flabellate = flabelliform,” things that are likely to mean the same thing such as “orbicular = round,” fine but unneeded distinctions like “pentagonal = polygonal,” and removal of subjective qualifiers (“sub-”, “usually”, “quite”). All of these behaviors are completely customizable.

Reading in the field controlled vocabularies leads to what is referred to here as “column-smashing” (Fig. 3b), where fields judged combinable are summarized according to their data type. At this stage, an additional spreadsheet representing taxa extracted by hand can be included, e.g., for data taken from direct measurements or small works such as single species descriptions that do not lend themselves to automated methods. Two types of output are produced: one with all of the data, and one with summarized single data points per cell. By default, the summaries are means for quantitative data, midpoints for count data, and random selections for categorical data. Finally, output can be filtered by missing data proportions, which functions primarily to exclude special descriptive material not shared among species. As well as outputting a raw harmonization of the data, the last function provided by the codes is a distance matrix that can be used to easily perform trait ordinations such as MDS (multidimensional scaling). The distance metric as described by Folk et al. (2019) is a hybrid metric comparable to Gower’s distance, partitioning categorical and quantitative data separately but with the ability to specify arbitrary weights to each.

*Ground-truthing—*To capture Type I (false-positive) and Type II (false-negative) errors, which respectively measure incorrectly captured and missed trait variation, we manually scored the extracted trait data using interactive HTML outputs. The two measures were calculated slightly differently: Type I error was calculated with the denominator as the complete count of partial traits since these are reported exhaustively by FloraTraiter and could be individually checked. Type II error was instead calculated with the denominator defined as the count of full traits (i.e., counting whether content was captured for each basic trait topic, while not scoring how verbal details were parsed out) because it was not possible to manually duplicate the tokenization process. As a manageable use case involving diverse plants with many specialized structures, we focused on all available species-level treatments for Betulaceae in *Flora of North America* and *Flora of China* as represented on *eFloras*, comprising 125 species.

*Worked example—*Fagales treatments as a whole were used, again sourcing from *Flora of North America* and *Flora of China*, to demonstrate one typical downstream use focused on quantifying trait spaces. Using the MDS procedure noted above, which is well-adapted to sparse matrices, we quantified trait spaces across Fagales using all morphological features populated with at least 5% of species. This analysis also demonstrates the reconciliation of automated and user-coded features, which are demonstrated in the empirical GitHub repository (https://github.com/ryanafolk/fagales_traits) and were scored from additional online sources including the *Flora of Australia* (https://profiles.ala.org.au/opus/foa) and *Flora Malesiana* (https://portal.cybertaxonomy.org/flora-<u>malesiana/</u>). This repository demonstrates the original extraction and conversion to controlled fields.

*Codebase availability—*The codebase for FloraTraiter is disseminated on GitHub at https://github.com/rafelafrance/FloraTraiter. A worked example processing Fagales content from *eFloras* is deposited at https://github.com/ryanafolk/fagales_traits.

## RESULTS

*Parsing success—*FloraTraiter was implemented on the complete Fagales treatments on *eFloras* as represented in *Flora of North America* (212 taxa successfully extracted) and *Flora of China* (518 taxa extracted). On average 55 traits per species were captured. The most densely populated traits included leaf shape (83.0%), leaf size (79.6%), leaf margin (61.0%) and surface (54.5%).

*Ground-truthing—*Type I (false positive) error rates were under 1% (0.98%; Fig. 5a and 5c break this down by source and genus). Many errors tend to be removed when filtering on missing data because they often involve uncommon or very detailed traits. This was because Type I errors were generally of the linkage type; a measurement was captured and parsed correctly but associated with a wrong, spatially adjacent trait. Most often such association issues happen for less common traits such as vein numbers (accounting for almost all errors; this is reflected in the higher error in *Flora of China*; Fig. 5a) that were unparsed as such; during the iterative process summarized in Fig. 2 unused tokens were then likely reused and misassigned. This example is illustrative of the most straightforward way to address Type I error: maximizing the text content that is parsed and explicitly assigned to traits. Rare traits are also where we see the most Type II errors, which was about 4.7% (i.e., the average treatment captured 95.3% of traits). Some of this was due to phenological traits (flowering before leaf-out, etc.), scent and taste descriptors, and other aspects that we decided not to capture because they are not basic morphology; many more were detailed treatments of taxon-specific topics like extensive prose on bark structure, winter buds, and elaborations of fruit and seeds that did not contain straightforward scorings. This last point is likely responsible for the higher Type II error in *Flora of North America* (Fig. 5b); there is a greater proportion of less structured, informal discussion of traits in this source. Connected to this, invariant “nuisance” traits were sometimes captured because the original description mentioned structures without providing relevant details. These were not counted as errors since they are uninteresting rather than incorrect and can easily be removed. For instance, lateral veins were captured as present with no detail because they were mentioned in relation to pubescence or other features. The capture was also sometimes incomplete because minor details were not included (this was generally not recorded as Type-II error because partial traits were not counted). Examples include color change over time or extensive prose on hair or ridge distributions. Excluding these from consideration is reasonable because these are unlikely to be useful in a morphological analysis without further standardization.

**Fig. 5.**
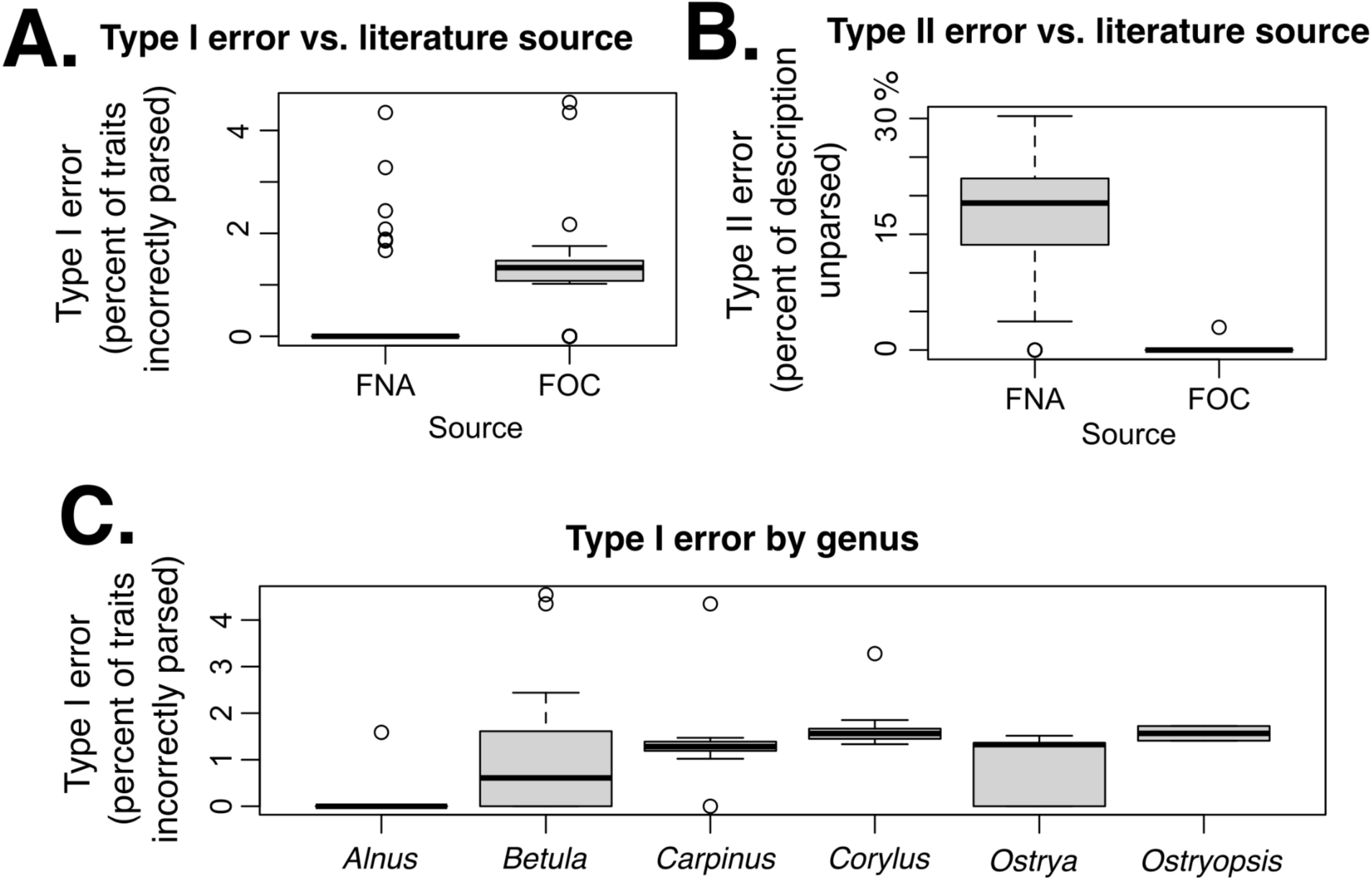
False positives (Type I error) and false negatives (Type II error). Type I error is shown vs. the two flora sources (A) and genus (C). Type II error is shown vs. flora source. Much of the structuring of errors appears to reflect differing editorial practices. Type II error was higher in the *Flora of North America* (FNA) treatment; the missing portions largely comprised prolix discussions of fine features. Type I error was higher in *Flora of China* (FOC), reflecting extensive discussion of leaf vein characters that were not parsed in detail but could easily be discarded.

*Worked example—*The MDS analysis (Fig. 6a; each dot represents a taxon) demonstrated differences in morphological variability among families, with Fagaceae (the most species-rich family) showing a wide spread and Betulaceae showing the smallest morphospace as quantified by between-species morphological distances. The within-family results show differing levels of morphospace occupancy per family, with Fagaceae showing the strongest variability; it alone occupies more than half of MDS axis 1 (Fig. 6a), which largely captures variation in vegetative and reproductive size (below) and therefore reflects morphological diversification in this family. Between-family comparisons likely underestimate morphospace divergence due to lack of homology; Casuarinaceae in particular possesses numerous fields not shared with close relatives relating to its specialized morphology. We note in this connection that this challenge is not unique to automated analysis; manual extractions possessed similar patterns of unshared fields between divergent taxa.

**Fig. 6.**
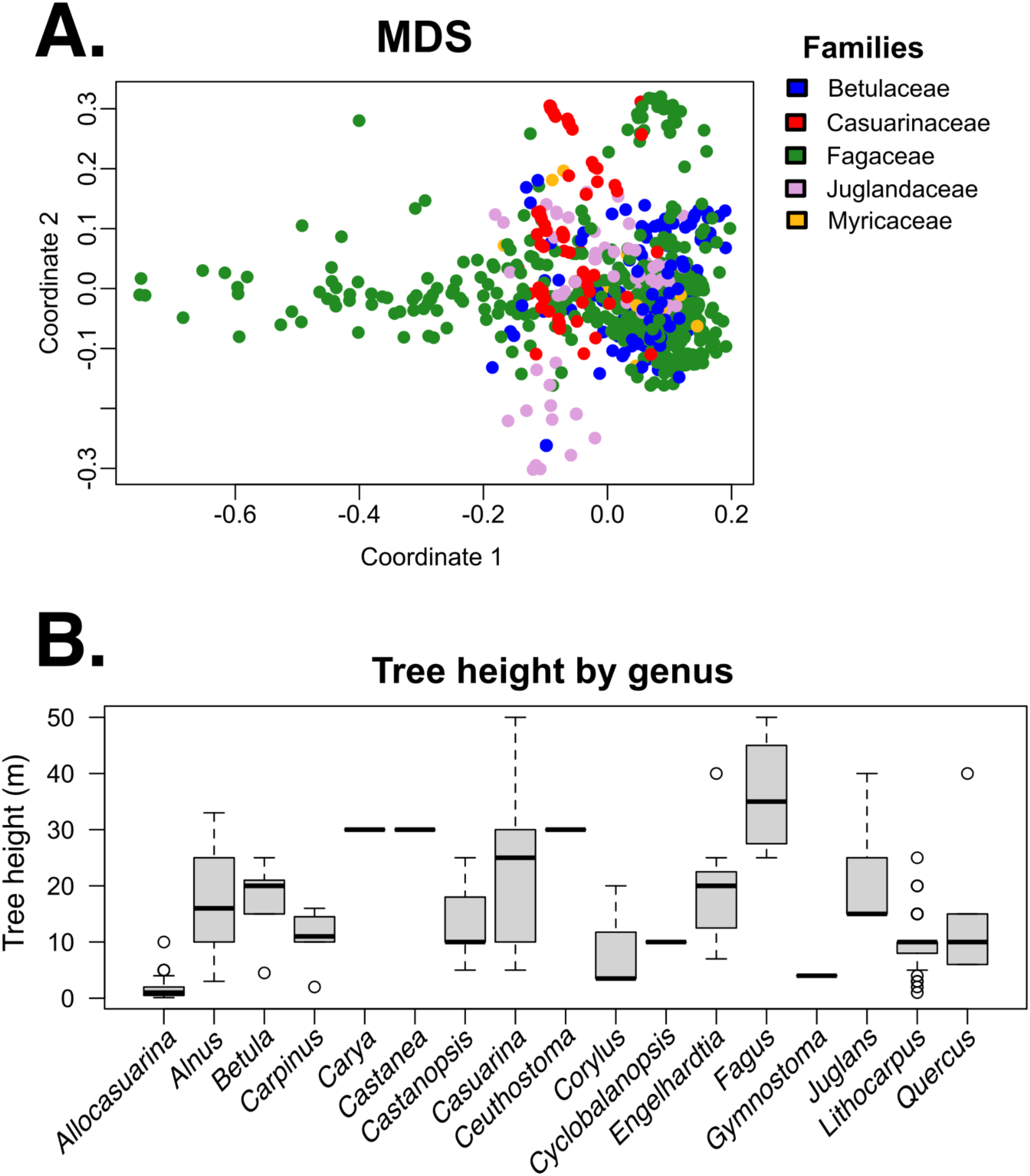
Worked example in Fagales. In this proof-of-concept, sampling primarily reflects taxa available in standard online floras; note Nothofagaceae is missing. (A) MDS, demonstrating variability of overall morphospaces within and between families. Each dot represents one taxon (mostly species; subspecific and higher-level taxa are also plotted). (B) Tree height plotted by genus.

Following Folk et al. (2019), “loadings” in the MDS analysis were evaluated by univariate *R*^2^, which quantifies variance explained by each ordinated axis. In the first axis, the most important traits were leaf width (*R*^2^ = 0.85), nut width (0.74), leaf length (0.72), and nut length (0.54); hence overall axis 1 captures size. The most important traits in the second axis were leaf duration (0.81), branch shape (0.28; this trait mostly captured information related to position such as “ascending”), peduncle shape (0.23), and inflorescence shape (0.20); hence overall axis 2 represents shape descriptors. Finally, Fig. 6b demonstrates an individual trait, showing tree height (the typical quantification of body size in plant ecology) against genera with sufficient subspecific sampling.

## DISCUSSION

A lack of trait data is a major obstacle in plant science because traits underlie how organisms interact with their environment and other organisms. However, researchers are met with the enormous challenge of collecting many traits from many taxa. The effort needed, for example, at the familial or ordinal level for many speciose clades, is immense, especially if starting from scratch. For many of the most basic traits, the problem is not that the data do not exist but that researchers cannot easily assemble them. With resources like the *Biodiversity Heritage Library* (https://www.biodiversitylibrary.org/) and the *Botanicus* project (http://www.botanicus.org/), large troves of trait data are available but disseminated in a way that currently lend themselves only to specific subfields and applications. As we have argued, unlocking the potential of biodiversity literature will be a key component of overcoming the lack of trait data, in a complementary role with other approaches such as generating new in-situ field-based measurements. Yet this heritage data is vast in scope and can be dissimilar in content, which highlights the need for automated and semi-automated approaches, especially since these data can not only be used once and for one purpose, but also re-used by others in new applications as befits their value. Such FAIR data re-use approaches (Wilkinson et al., 2016) have not been particularly well-facilitated by traditional scientific literature dissemination modalities.

Here we developed and demonstrated the application of FloraTraiter, a fully open-source natural language processing approach to extracting computable trait data from taxon descriptions in floras, monographs, and similar biodiversity literature. FloraTraiter is close to full automation, but contains several human loops that add customizability as well as quality control. Some human roles are very specific to particular projects, such as overcoming challenges in digitizing treatments and in controlling and interpreting output structures. But most centrally, expert botanist perspectives are built into the NLP model, a key especially for specialized contents where applying previously developed approaches “out of the box” may result in limited parsing (Endara et al., 2018). We identify a number of challenges related to the structure (or lack thereof) with the data that will inform future efforts (see also Supplement S1), but also demonstrate an accuracy already sufficient for re-use in diverse applications and in a form that will facilitate further curation.

FloraTraiter fits into recent efforts unlocking similar literature data in other organisms, such as lepidopterans (Shirey et al., 2022) and vertebrates (Guralnick et al., 2016). The present effort differs from Shirey et al. (2022) in that much of that work relied on human effort to assemble and parse text blocks, with automation for only the most simple of continuous traits (e.g. wing length). Unlike Guralnick et al. (2016), floral traiter interrogates much more complicated written text blocks, and assembles a much more complicated set of both continuous and categorical traits. Our work not only shows the power of semi-automation to scale up trait assembly, but also brings home the critical importance of a more explicit model for collaboration between botanists and programmers to structure improving language processing models. Similar efforts have also been recorded in plants (e.g., Endara et al., 2018 report a previous NLP approach), but we have moved here beyond proof-of-concept in a large scope with a dataset comprising hundreds of taxa ready for reuse.

*Challenges encountered—*Challenges and tips in the effort overall are summarized in Supplement S1. Here we will focus on empirical properties of the output. Type I error rates are low (<1%) and similar to those reported in previous NLP efforts (Endara et al., 2018). The specific Type I errors found were attributed to the completeness of parsing, with information involving unparsed traits most prone to incorrect trait linkage. This suggests a straightforward strategy in addressing Type I error focused on maximizing the vocabulary that would enable tokenization in the first steps of FloraTraiter. Type II error was also fairly low (< 5%); this is more difficult to control as it was found to be highly dependent on authorial style among sources, as standard taxonomic styling lends itself to NLP but more prolix approaches to the discussion of traits tend not to be parsed. However, excluded trait information tends to be highly detailed and may not be specifically of interest for many projects. In Betulaceae, this often involved relative flower and leaf phenologies (not included by decision) and extensive bark descriptions (which could be challenging to score even for a human observer).

As noted by Endara et al. (2018), matrices created by automated trait extraction tend to be sparse: basic leaf and other traits are well filled-in, but other traits less so. This is a feature rather than a bug; it reflects the higher thoroughness of an automated approach as opposed to the more selective strategy a human observer would likely take. Trait presence is primarily a function of both morphological specialization and editorial differences among sources. At a large taxonomic scope, fields must be invoked to cover all extracted traits, but within subtaxa taxonomists generally will focus only on those traits that differ between species. Some traits are also not straightforward to homologize because features are not shared across taxa in a straightforward way (for instance, when comparing details of berries to samaras, or when comparing Casuarinaceae overall to other families). In the Betulaceae application used for error analysis, there there were fewer problems because Betulaceae trees are fairly similar, but inflorescence structure is diverse and comparing them across genera requires enforcing controlled fields and synonymies.

*Conclusions—*Among the extracted traits that are broadly scored across species are body size, measurements and shape descriptors of leaves and flowering and fruiting structures, scorings of plant sex, and other attributes that have broad use potential. For instance, these traits capture aspects of plant and leaf size that have recently seen global-scale studies in a phylogenetic (Testo and Sundue, 2018) or spatial framework (Baird et al., 2021). Aside from investigations of targeted questions, traits could be used as a whole to capture regional or site-level morphospaces, e.g. via calculation of convex hulls (Cornwell et al., 2006) or other methods; ordinations of morphospaces could also be used in a phylogenetic context to investigate trait evolution (Folk et al., 2019). These examples underline the value of making biodiversity literature more available and computable (Thessen et al., 2012; Folk and Siniscalchi, 2021) to broad audiences beyond the traditional readership of such literature, including scientists who will be able to identify important, unusual, and creative applications. Most centrally, FloraTraiter demonstrates the major promise of combining expert perspectives with guided automated approaches, and the promise of tools designed to facilitate curation for non-specialists. Future challenges will include data curation and reconciliation challenges, and assuring that outputs are broadly usable for the long term and conform to community definitions. We also foresee that large language models and model-based parsing may rapidly improve and provide an alternative to the rule-based parsing shown here; still those models are likely to require the same set of lexicons that are specific to botany and therefore to benefit from collaboration with domain experts. Here we show a pathway to making data and methods available that can serve as an important first step towards broadly available plant trait data for the community.

## ACKNOWLEDGMENTS

We first thank the collective efforts of taxonomists over hundreds of years; the automated approach focuses on recent works but this still substantial legacy depends on the expertise of botanical experts. For input on our work we thank Leonardo Borges, Yago Barros, Carolina Siniscalchi, and Joshua Doby. This work was supported by NSF DEB-1916632.

## Supplement S1. Guide to generalizing NLP and overcoming challenges

This supplement documents insights the authors had in preparing FloraTraits for diverse literature sources, as well as more general challenges for those new to complex text parsing. The advice in this document is not general to NLP however; it is aimed at those parsing complex and jargon-laden text full of unusual abbreviations and other field-specific customs that confound prebuilt language models. In this connection, it should be noted that FloraTraiter parsers are geared specifically towards information extraction of natural history documents and notes, and therefore may not be generalizable to non-natural history materials and may require customization based on organism. For instance, ‘762-292-121-76 2435.0g’ has a very specific meaning to people who work with smaller vertebrates, and ‘(12-)23-34 × 45-56 cm’ is obviously length and width size range measurements containing a minimum length to people who work with plant treatments.

## Technical Hints and Challenges

**Hint: An exposition (not explosion) of terms**. The aim of FloraTraiter in terms of NLP is information extraction (**IE**). We are finding traits using phrases and rules, linking them using other rules, and then extracting information from them in callback functions. The established term Named Entity Recognition (**NER**) refers in this context to finding the traits, and Named Entity Linking (**NEL**) refers to linking the traits.

**Hint: Use a package like spaCy** from the start. There are other options for NLP; in the Python world Stanza and NLTK are additional options. spaCy was identified as best based on initial tests in our data. In implementing initial tests, consider that the main purpose of using these packages is to off-load as much of the parsing work to computational experts as possible. Writing parsers for this kind of data is hard enough without a strong foundation.

spaCy is pipeline-based and allows a user to swap in or out pipeline segments as needed. This functionality is essential because our parsing needs are unique and pertain to various steps in the process. As an additional consideration, the spaCy team is responsive, patient, and very helpful.

**Hint:** Adding to the above hint, it is best to **leverage as much of spaCy as possible**. spaCy has a default pipeline. While some steps must be customized and the others removed, the emphasis is to try to use as much as possible directly from spaCy. FloraTraiter in this case specifically uses a modified tokenizer, the default Parts Of Speech (POS) model, and the default lemmatizer (the portion of that model that determines the lemma). In certain cases spaCy’s default NER model was used as input to custom rules.

**Challenge:** A note of caution about the hint above: **use models not of your making cautiously**. Most of what the spaCy team produces is expertly implemented but some of the models will change when there is a minor revision, leading to problems with custom content. For instance, a model was implemented on sentences to do a dependency parse on each one. Everything worked well until there was a minor spaCy upgrade and all of the dependency parses changed slightly, leading to all rules breaking. By contrast, the POS model has changed but not in any way that affects the rules implemented here.

**Hint:** The basic pipeline in FloraTraiter is:

1. Use a modified tokenizer to split the input text.
2. Allow spaCy to implement its methods on parts of speech, lemmas, and named entities.
3. For every trait I do some variation on the following:
  a. Find words and phrases (terms) to use as an anchor for the trait. Also find words and phrases that will never be in the trait and use them to block out noise. Sometimes these terms are the final trait.
  b. Find patterns of words and symbols around those terms that build a trait. This step is often repeated to continue to build up larger and more complicated traits.
  c. For each trait found, parse its details in a callback function.
  d. Next, cleanup any partial traits from above so that leftover words can be used in other traits.
4. Traits are linked with other traits. For instance a size may refer to a leaf size, flower size, or a plant size.

**Hint:** The spaCy tokenizer was originally built to handle web text like Wikipedia, twitter tweets, etc. It was not built to handle the idiosyncratic text found in specialist sources. We find that **customizing the tokenizer** by removing problematic character sequences and splitting on more characters helps substantially.

**Hint: Terms are words or phrases**. It is important to incorporate as many of these as possible from domain experts such as expert botanists. FloraTraiter uses one or more spaCy phrase matchers to find these; it is fast and can build and find millions of terms (like those from a taxonomy or gazetteer) in seconds.

**Hint: Rules are patterns of tokens** and are analogous to regular expressions for tokens. Rules can get complicated but what FloraTraits does to reduce this complexity is to build up traits in stages. This can greatly reduce the number of trait permutations to search. For instance, to build up a taxon trait (in simplified terms), FloraTraiter first:

1. Looks for all binomial, monomial, and rank terms in a document:
  a. Binomial: “Genus species”
  b. Monomial: “FamilyName” etc.
  c. Rank: “subspecies”
2. Then it searches for species patterns like “binomial” or a binomial followed by a lower rank and monomial like “Genus species subsp. subspecies” and gives it a taxon label.
3. Next it sees if there is an authority attached to the taxon from step 2 by looking for a taxon followed by a name like “taxon (name) name”. This will still have the taxon label.
4. After that it looks for an extension lowering the taxon’s rank. Like: “taxon rank lowerMonomial (name)”. This extends the taxon and lowers its rank.
5. Finally it removes any unused binomials, monomials, & ranks so they can be used in other traits. There are already well over 100 patterns for parsing taxons, so trying to do all of steps 2 through 4 in a single set of patterns would grow the numbers of patterns exponentially.

Hint: A trait in FloraTraiter is nothing more than an annotated spaCy entity which is itself nothing more than an annotated spaCy span.

**Hint:** You can **strategically order the traits** to simplify trait parsing. Simple strategies include parsing latitude and longitude before looking for size traits. This makes sure that sizes do not interfere with lat/longs and lat/longs are guaranteed to not interfere with sizes because lat/longs are either labeled or in a specific format.

**Hint:** You can **allow some traits to overwrite other ones**. This allows you to disambiguate traits by context. For instance we can identify a color like “Brown” and later on if it is next to a taxon change it to an authority or if it is next to a “county” label change it to a county.

**Hint:** Close inspection of the FloraTraits rules will show that they do not look like the rules in the spaCy documentation. That is because FloraTraits **“compiles” the rules before they are used**. Experience demonstrates that, in looking at strings with descriptive labels vs. JSON blobs, it is much easier to reason about what a patterns is doing and why it may or may not contain a bug with the former.

**Hint:** When there is a trait match (final or partial), FloraTraits parses its contents to **build up the trait data in a callback function**. This is done when the trait is matched because it is easier to parse a trait when it is separated into tokens. Sometimes traits are merged into a single token.

**Challenge:** The trait data is currently stored in a Python dictionary in FloraTraits. In the future it would be more effective to store **trait data in data classes**.

**Hint: Linking traits uses patterns** too. The patterns superficially look fairly simple, but there is some complicated code in the library module that does the actual linking. Under the hood, linking involves looking at all nearby traits and using a weighted count of the token distance between them to find the “closest” trait. The weights are mostly on punctuation so that it costs more to cross a period than a semicolon and a semicolon costs more than a comma. Some traits get linked multiple times (e.g., a flower could come in multiple colors) but other traits only get linked once (e.g., it does not make sense to have more than one flower size unless sexual dimorphism is involved). There is a fixed list of parent and child traits for linking.

**Hint:** A certain paranoia is justified about regression errors (in this context, code that is broken by external events such as package updates) and it is important to test accordingly. There are so many moving parts to the parsers that it is too difficult to consider them all when bugging. So, FloraTraiter uses **red/green testing**. When an error is encountered, the offending text is put into a test case, the test is run to make sure it fails, the code is fixed, and finally a rerun of the test ensures it passes. The tests can be accordingly numerous, and some of them repeatedly test code that is robust at this development stage, but the philosophy is to keep them until it is certain they are no longer needed.

## Collaboration Hints and Challenges

**Hint:** Get as much **starting data, terms & rules, as soon as possible**. This is the foundation for building the parsers and without this information they cannot be implemented effectively. This involves asking organismal experts very targeted and structured questions to obtain information, such as lists of trait synonyms.

**Hint:** Related to above, **get feedback** on the results as soon and as often as possible. Start parsing some results as soon as possible. Do not wait until all of the parsers are done; rather, get one parser done and send the results back to the experts for guidance. Put these results into an easy to digest format (in this case, colorized HTML reports, which organismal experts found very effective). Prebuilt tools are available for this purpose; spaCy module “displacy” provides much of the needed functionality.

**Challenge:** Parsing specialist text is **team work** and many errors in are the responsibility not of the programmer but of domain experts. Obvious errors are easily fixed, but subtle errors require deep engagement with the subject material.

**Hint:** There are **other people parsing** similar types of data that share their experiences online; YouTube videos are a particularly rich source of guidance. Two of the channels on YouTube that are relevant are the <u>spaCy</u> channel itself and <u>Python for the Digital Humanities</u>; these contain a mix of basics and deep exploration of specialist topics.

**Challenge:** There are genuine one-off scripts but parsing complex text never seems to be one of them. Do yourself a favor and **treat each parsing project like you’ll use it again**. This includes strong programming practices like documentation, testing, and modularity.

## Strategic Hints and Challenges

**Challenge:** Probably the biggest strategic hurdle in this project was **not prioritizing model-based parsing** soon enough in the project (see Introduction). Aside from rule-based parsing being tricky, labor-intensive, error prone, and tedious, it is also quite brittle. Most rules will fail or worse, yield an incorrect parse, when presented with a new pattern, whereas models can sometimes adapt. Both models and rules suffer from “model drift” or “data drift” (a decay in predictive power as the underlying processes generating the data change) but for rules it can be catastrophic. Errors include turning dates and ID numbers into counts and administrative units into names, etc. In the long run, models are less work.

However, given the nature of what is being parsed in specialist text, **some rule-based parsing** is both inevitable and practical. Using rules to bootstrap model training data is helpful. Additionally, a hybrid parser that uses both a model and rules is a powerful parsing technique overall.

The **models can be simple**. There are cases of Hidden Markov Models used effectively for NER. What FloraTraiter does is unique in detail but older techniques may also perform well; a simple NER model was tested preliminarily on scientific prose with excellent results.

**Challenge:** Another major hurdle was awareness of **free NLP packages** and techniques readily available, leading to unnecessary effort to develop everything in-house. This puts responsibility on the developers to implement challenging mechanistic details that are already freely available from experts in robust packages.

